# Vasomotor Pulsation Becomes a Driver of Cerebrospinal Fluid and Electrophysiological Dynamics in Sleeping Human Brain

**DOI:** 10.1101/2024.10.16.618430

**Authors:** Tommi Väyrynen, Johanna Tuunanen, Heta Helakari, Ahmed Elabasy, Vesa Korhonen, Niko Huotari, Johanna Piispala, Mika Kallio, Maiken Nedergaard, Vesa Kiviniemi

**Author notes:** **Correspondence to:** Tommi Väyrynen.

## Abstract

Sleep is essential for maintaining brain tissue homeostasis, which is facilitated by enhanced cerebrospinal fluid (CSF) solute transport. Infra-slow (<0.1 Hz) vasomotion, CSF flow, and electrophysiological potential all increase during sleep, but their contributions as potential drivers of CSF flow in human brain remain unknown. To investigate this, we recorded the three signals in healthy volunteers across sleep-wake states using 10 Hz functional magnetic resonance imaging (fMRI BOLD), electroencephalography (DC-EEG), and functional near-infrared spectroscopy (fNIRS). We then analyzed the directed coupling patterns using phase transfer entropy (TE).

In the awake state, electrophysiological potential and water concentration changes both predicted vasomotor waves across the brain, possibly reflecting functional hyperemia. During sleep, this coupling reversed, with vasomotor waves instead predicting electrical changes and CSF flow in cortical areas. Furthermore, we found that the amplitude of these dynamics increased during sleep, highlighting the critical role of physiological oscillations in sleep-associated brain fluid flow.

## Introduction

Sleep induces infra-slow oscillations (<0.1 Hz) in the cerebrospinal fluid (CSF), which facilitate the clearance of accumulated solutes and metabolites from the cerebrocortical parenchyma^1,2^. These oscillations result from low-concentration norepinephrine (NE) waves originating from the locus coeruleus, which are known to induce vasomotor waves^3,4^. The slow vasomotor waves generate oscillations in cerebral blood volume, which directly reflect upon CSF flow, consistent with the Monro-Kellie doctrine^4–6^. The NE-driven cortical vasomotion inflates the perivascular CSF space volume and thickness of astrocyte endfeet plastering the outer edge of perivascular spaces of the blood-brain barrier (BBB)^7–9^. Numerous human functional magnetic resonance imaging (fMRI) studies have demonstrated that vasomotor wave amplitude increases with sleep depth, suggesting a direct link between sleep and vasomotor activity^6,10–15^

Beyond its impact on vasomotor waves, sleep also enhances infra-slow oscillations in the direct current-coupled electroencephalogram (DC-EEG)^16,17^, which further modulate neuronal rhythms over a wide range of frequencies^17–19^. Among these rhythms, sigma power has a notable association with the declining phase of LC activity^1,20^ and with memory consolidation after sleep^21^. The infra-slow EEG oscillations are further connected to changes in cerebral blood volume (CBV), extracellular pH, [K+], and permeability across the BBB^22–24^. Recent works shows such synchronized neuronal activity to facilitate interstitial influx and efflux of electrolytes^25^. Synchronous EEG & fMRI studies have also linked infraslow, synchronized neurovascular activity to enhanced solute exchange between interstitial fluid and CSF during sleep^6,26^. The three types of infra-slow physiological oscillations - vasomotor waves, electrophysiological voltage shifts, and CSF flow – all increase in amplitude during sleep, yet the temporal and causal relationship among them remain largely unexplored, especially in humans.

A recent animal study showed that NE oscillations coordinate vasomotor waves, CSF flow, and EEG sigma power, and that sleep alters their correlation and lag structure^4^. However, most studies have examined these factors in isolation, thereby overlooking their potential mutual interactions. To identify the drivers of the sleep-induced increase in CSF flow and facilitated brain solute clearance in humans, we developed non-invasive multimodal imaging protocol devoid of artificial tracers. We thereby simultaneously measured the infra-slow vasomotor waves, electrophysiological DC-EEG, and CSF flow oscillations non-invasively from the human brain, using 10 Hz sampling of fMRI blood oxygen level dependent (BOLD) signal to avoid cardiorespiratory aliasing effects^27^. For CSF quantitation, we used a novel water-sensitized fNIRS technology^5,28^ recently shown to be crucial for sleep-induced glymphatic solute clearance^4^. Finally, to gain an understanding of the underlying causal patterns, we analyzed the interactions between these three signals using information theory-based phase transfer entropy (TE)^29^ across wakefulness and sleep states. Based on prior mice studies^1,4,9^ we tested the hypothesis that oscillations in the CSF water signal in human brain are driven by vasomotor waves during sleep.

## Results

To investigate the interconnected hemodynamic, water, and electrical oscillations in the human brain, we employed a multimodal neuroimaging setup (Fig. 1a). Our study included 24 healthy participants with an equal gender distribution (54% females) and a mean age of 25 years in both gender groups (Fig. 1b). Each participants completed two scanning sessions: one during wakefulness and another specifically targeting sleep. Experienced neurophysiologists (J.P., M.K.) manually classified sleep stages from the EEG recordings. From continuous sleep state-specific epochs across participants, we extracted in total 46 minutes of wakefulness, 40 minutes of NREM-1 sleep, and 28 minutes of NREM-2 sleep during active MRI scanning (Fig. 1c).

**Figure 1.**
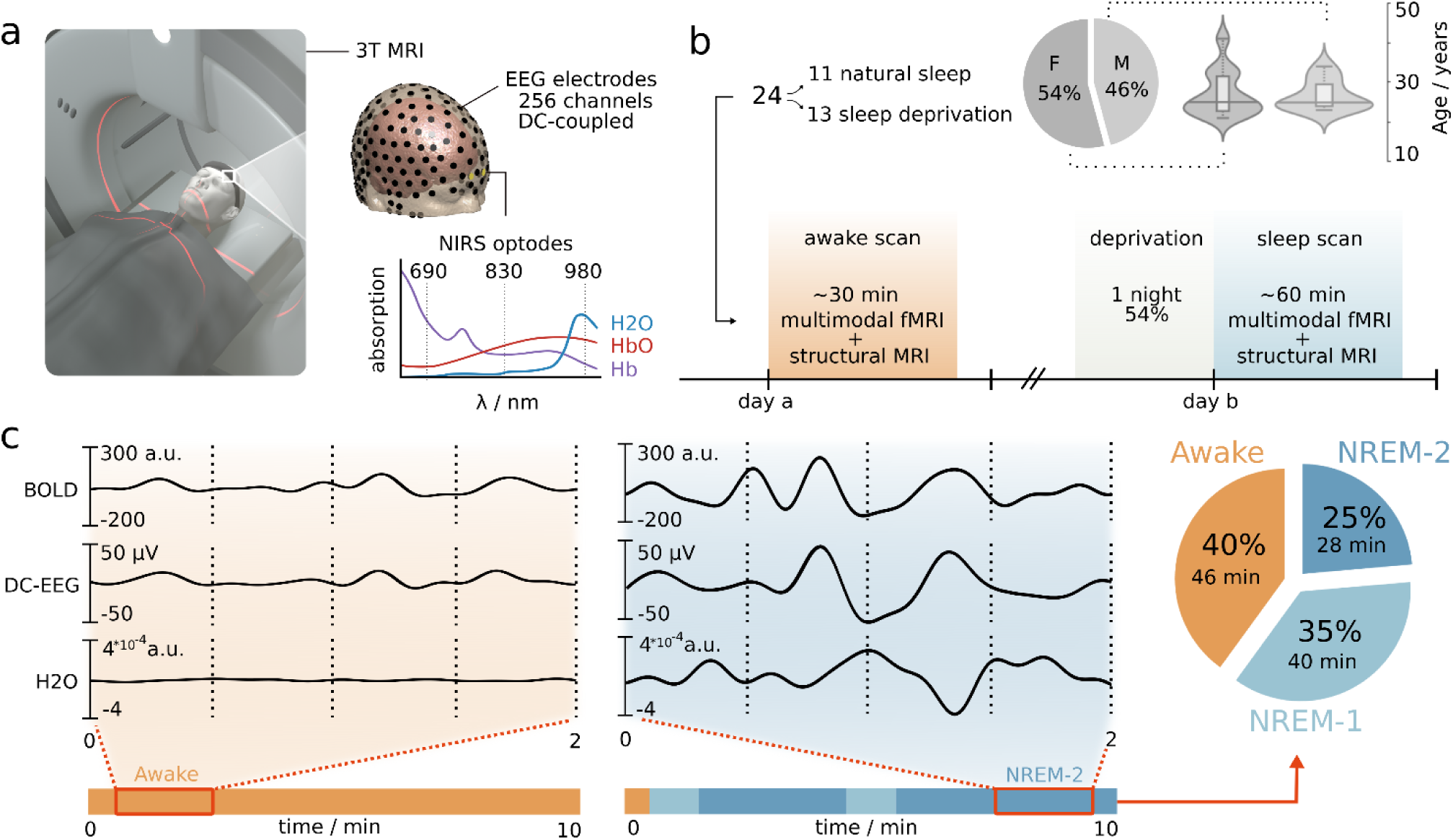
Multimodal imaging protocol for simultaneous measurement of brain water content, electrophysiological, and hemodynamic processes. (a) Measurements were conducted in a magnetic resonance imaging (MRI) scanner using a 10 Hz whole-brain magnetic resonance encephalography (MREG) sequence with simultaneous direct-current electroencephalography (DC-EEG) and functional near-infrared spectroscopy (fNIRS). We applied a water-specific NIRS wavelength (980 nm) to measure brain water concentration changes. (b) Participants underwent two scanning sessions comprising functional MREG and structural scans: one during wakefulness and another during sleep. The pie chart illustrates the gender composition of the (n=24) participants, while the violin plot depicts their age distribution. (c) Representative signals of infra-slow (<0.1 Hz) MREG, EEG, and water NIRS from one participant along with corresponding EEG-derived sleep scoring. For visualization, NIRS signals were calibrated with respect to MREG. The pie chart shows the proportions of measurements across three arousal states (awake, NREM-1 and NREM-2) after searching for continuous segments.

### Increased infra-slow (<0.1 Hz) hemodynamic and electrical oscillations in sleep

We first examined the behaviors of different modalities in the frequency domain to confirm our expectation of increased power across all modalities during sleep. Using two-minute epochs of verified wakefulness and sleep (NREM-1/2), we computed the time-frequency estimates (Fig. 2a) within the 0.01 to 5 Hz frequency range. Visual inspection confirmed the presence of three distinct frequency bands: infra-slow, respiratory, and cardiac ranges.

**Figure 2.**
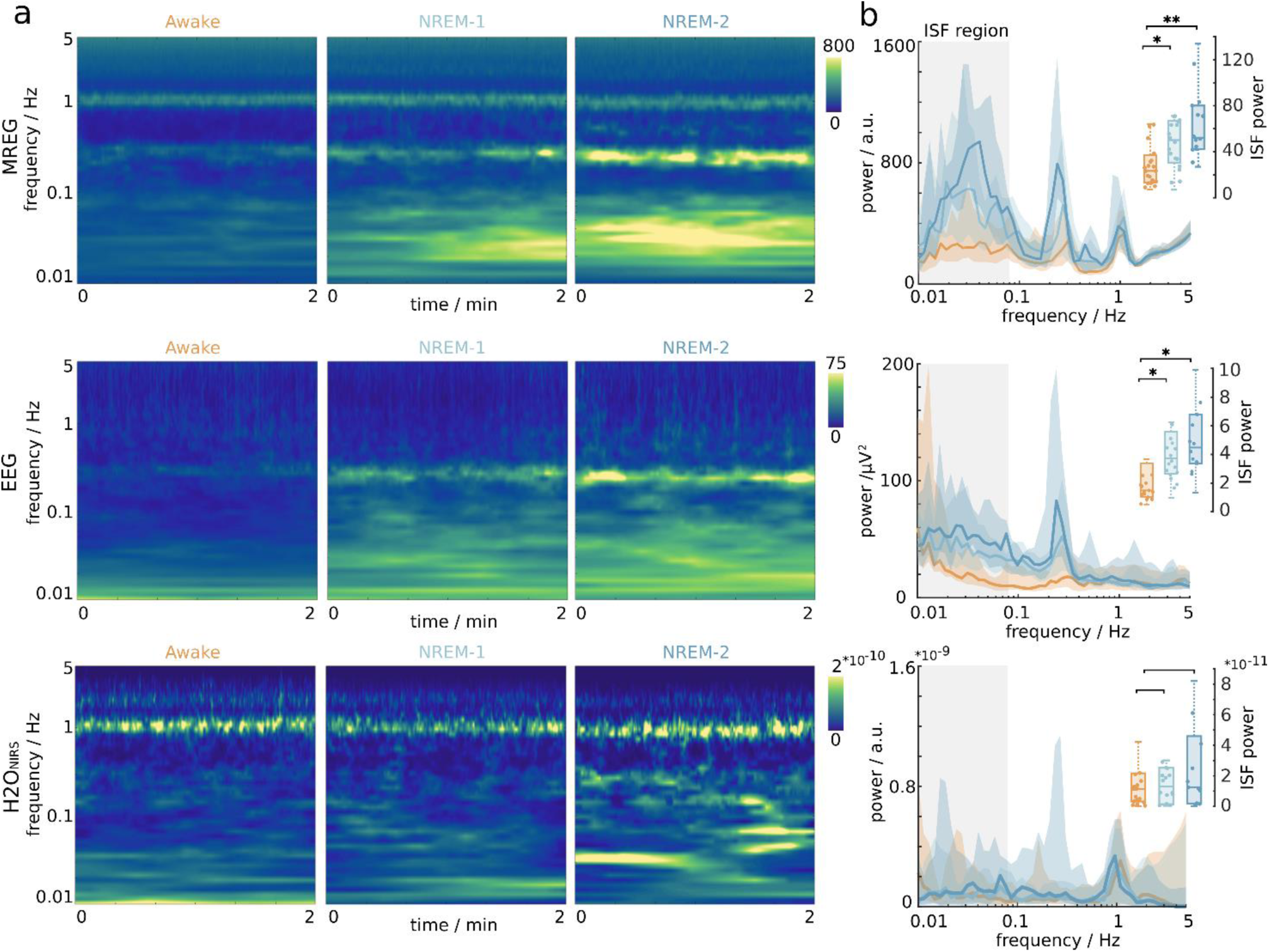
Sleep increases infra-slow power (<0.1 Hz) of magnetic resonance encephalography (MREG) and electroencephalography (EEG). (a) Time-frequency estimates illustrate the average spectral powers for MREG, EEG, and functional near-infrared spectroscopy (NIRS) within sleep state-specific epochs. (b) Time-collapsed power spectra depict frequency as a function of power, where the infra-slow frequency range (0.01-0.08 Hz) serves for statistical comparison. Data points corresponding to wakefulness are denoted by orange, and non-rapid eye movement sleep by light blue (NREM-1) and dark blue (NREM-2). The solid lines represent the median, while the shaded area indicates the middle 50% of data points. Statistical significance is denoted by asterisks (*padj*< 0.05*, 0.01**, 0.001***). The cardiac 1 Hz signal is missing from the EEG spectrogram due to fMRI related cardioballistic artifact removal.

In the transition to sleep, there was a clear increase in the infra-slow power of MREG signal (Fig. 2b), peaking at a frequency of 0.03 Hz. The group mean infra-slow power rose from *PA*=23.3 (IQR: 23.6) during wakefulness to *PN1*=50.1 (IQR: 36.3) and *PN2*=52.2 (IQR: 37.8) during sleep (A-N1: *Z(23,20)*=-2.59, *padj*<0.05*; A-N2: *Z(23,14)*=-3.46, *padj*<0.01**). We observed a similar trendin EEG signals, with the infra-slow power increasing from *PA*= 1.49 (IQR: 2.34) during wakefulness to *PN1*=3.73 (IQR: 2.93) and *PN2*=4.48 (IQR: 3.40) during sleep (A-N1: *Z(23,20)*= −2.42, *padj*<0.05*; A-N2: *Z(23,14)*= −2.61, *padj*<0.05*). However, there were no corresponding changes in the NIRS power measurements across arousal states, with values of *PA*= 1.15 (IQR: 1.84)*10^-11^, *PN1*=1.29 (IQR: 2.38)*10^-11^, *PN2*=1.24 (IQR: 4.40)*10^-11^. Consistent with our expectations and earlier results, power in the infra-slow range increased significantly in MREG and EEG measurements in proportion to sleep stage^12,17^.

### Sleep-induced local vasomotor drive of brain’s infra-slow electrical and water oscillations

Based on recent observations in mice^4^, we hypothesized that infra-slow vasomotion (BOLDMREG) could be linked to electrophysiological DC-EEG and macroscopic water fluctuations (H2ONIRS). To reveal the coupling patterns, we measured the information transfer among these signals using phase transfer entropy (TE) in bits (Fig. 3a), where TE = 0 indicates no connection between the phase time series. During wakefulness, we found that electrophysiological EEG changes predicted the slow vasomotor waves of the BOLDMREG signal (Fig. 3b). Furthermore, the H2ONIRS phase transitions also predicted the BOLD signal (Fig. 3d, shown in blue colors) across the entire brain, as indicated by a coherent negative net information transfer. In both NREM sleep states, the coupling patterns reversed locally, such that BOLDMREG started to predict the EEG and H2OEEG changes within specific cortical areas. This reversal is visualized as yellow-red regions against an otherwise blue brain map (Fig. 3b,c). The positive net information transfer was concentrated in parasagittal cortical regions, including the frontal, parietal, sensorimotor, and temporal areas.

**Figure 3.**
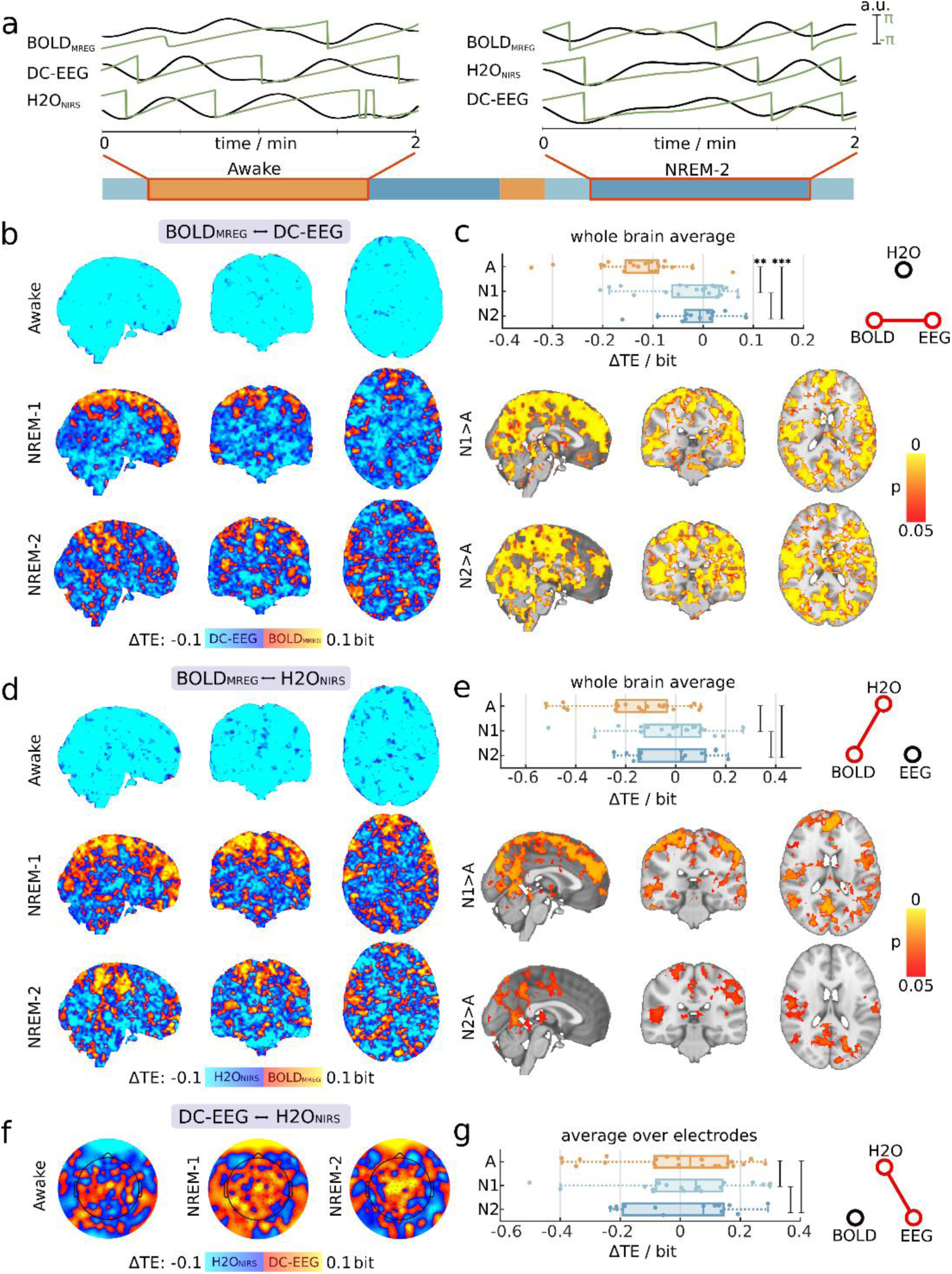
Sleep-induced local reversals in coupling patterns. (a) We utilized the phase time-series (green) of infra-slow signals (black) within sleep state-specific epochs. Transfer entropy in the phase domain (TE) was used to calculate information transfer. The sign of *ΔTE* indicates the net direction of information transfer, as also highlighted in the color bars. (b) Blood oxygen level-dependent (BOLDMREG) and electroencephalography (EEG) coupling patterns were assessed in wakefulness and non-rapid eye movement (NREM)-1/2 sleep. Positive values correspond to BOLDMREG prediction, while negative values indicate the net prediction of EEG. (c) We calculated the average TE over the whole-brain volume, where asterisks denote significance of differences (*padj*< 0.05*, 0.01**, 0.001***). Below, the spatial distributions of statistically significant areas (*padj*<0.05) are shown in comparisons to wakefulness. (d) Similarly, we present TE between BOLDMREG and cortical water dynamics (H2ONIRS), where positive values indicate BOLDMREG prediction, and negative values indicate prediction of H2ONIRS. (e) Whole-brain averages for BOLDMREG↔H2ONIRS coupling and spatial distribution of significant differences. f) Topographies show coupling patterns between EEG and H2ONIRS and g) average TE taken over electrode space.

The reversed information transfer of BOLDMREG↔EEG coupling during NREM sleep was prominent enough to appear in whole-brain averages (Fig. 3c). The TE values shifted from *ΔTEA=*-0.11 bit *(IQR:* 0.07 bit) during wakefulness to *ΔTEN1=*1.8*10^-3^ bit (*IQR=*0.09 bit) during NREM-1 and *ΔTEN2=*-0.01 bit *(IQR=*0.05 bit) during NREM-2. These shifts reflected statistically significant increases (A-N1: *Z*23,20=-3.4, *padj*<0.01**; A-N2: *Z*23,13=-3.7, *padj*<0.001***; N1-N2: *Z*20,13=-0.06, *padj*=0.96). Similarly, BOLDMREG↔H2ONIRS coupling demonstrated an increasing trend over the whole-brain averages (Fig. 3e), without reaching significance.

The median information transfer was *ΔTEA=*-0.12 bit (*IQR=*0.20 bit) during wakefulness, which increased to *ΔTEN1=*0.02 bit (*IQR=*0.22 bit) during NREM-1 and *ΔTEN2=*0.27 bit (*IQR=*0.02 bit) during NREM-2 sleep. These cortically dominant changes were not significant for the whole brain averages: (A-N1: *Z*22,20=-2.4, *padj*=0.06; A-N2: *Z*22,13=-1.6, *padj*=0.18; N1-N2: *Z*20,13=0.4, *padj*=0.87).

BOLDMREG→EEG predictions spanned a wider cortical area (*padj*<0.05) than the smaller, but overlapping BOLDMREG→H2ONIRS prediction (Fig. 3c, e bottom). Within these reversed prediction areas, 81% of voxels showing a BOLDMREG→H2ONIRS prediction increase in NREM-1 also showed a significant increase of BOLDMREG→EEG, whereas there was 98% overlap for NREM-2 sleep. Notably, most subcortical brain regions continued to exhibit negative prediction during sleep, where EEG and H2ONIRS phases continued to predict BOLD vasomotion.

We further examined the last candidate of interaction pairs: EEG↔H2ONIRS. Here, the topographies demonstrated more stable coupling patterns with respect to arousal state as compared to the other interactions noted above (Fig. 3f). Generally, the central upper vertex electrodes showed continuous predictive drive of EEG over H2ONIRS, while more peripheral, distal electrodes presented an opposite prediction: H2ONIRS→EEG. The average *ΔTE* over all electrodes showed that the prediction EEG→H2ONIRS was predominant, without significant alterations with respect to arousal state (Fig. 3e): (A-N1: *Z*22,20=-0.14, *padj*=0.9; A-N2: *Z*22,13=-0.29, *padj*=0.9; N1-N2: *Z*20,13=0.13, *padj*=0.9).

In line with our hypothesis, we found directional coupling among infra-slow brain hemodynamics, water and electrical changes across arousal states. Wakefulness was found to be a coherent state, where water dynamics and electrophysiological brain changes both predicted BOLD vasomotion. In NREM sleep, we found two coexisting mechanisms across brain: BOLD vasomotion coordinated the interactions in cortical structures, while deeper brain structures continued to exhibit dynamics similar to that in the awake state.

### Speed and power of the vasomotor waves co-localize with reversed CSF drive in primary sensory cortices

We have previously shown with MREG BOLD scans that pulsation power and speed of infra-slow vasomotor brain waves both increase during NREM-sleep^12,30^. Here, we further explored whether these changes in vasomotor pulsation characteristics are associated with the same brain regions where the sleep related vasomotor drive occurs. The increased power and speed of BOLD vasomotion were indeed present in the same brain regions where vasomotion predicts water and electrophysiological oscillations, especially in the primary sensory brain regions, upper cerebellum, thalamus and posterior insula (Fig. 4a & Supplementary Fig. 1). The spatial overlap of the increased vasomotor propagation speed with the areas of reversed electro/hydrodynamic CSF drive suggests that (peri)venous flow is either less restricted in these areas, or that the increased vasomotor power imposes a faster flow through the porous brain tissue.

**Figure 4.**
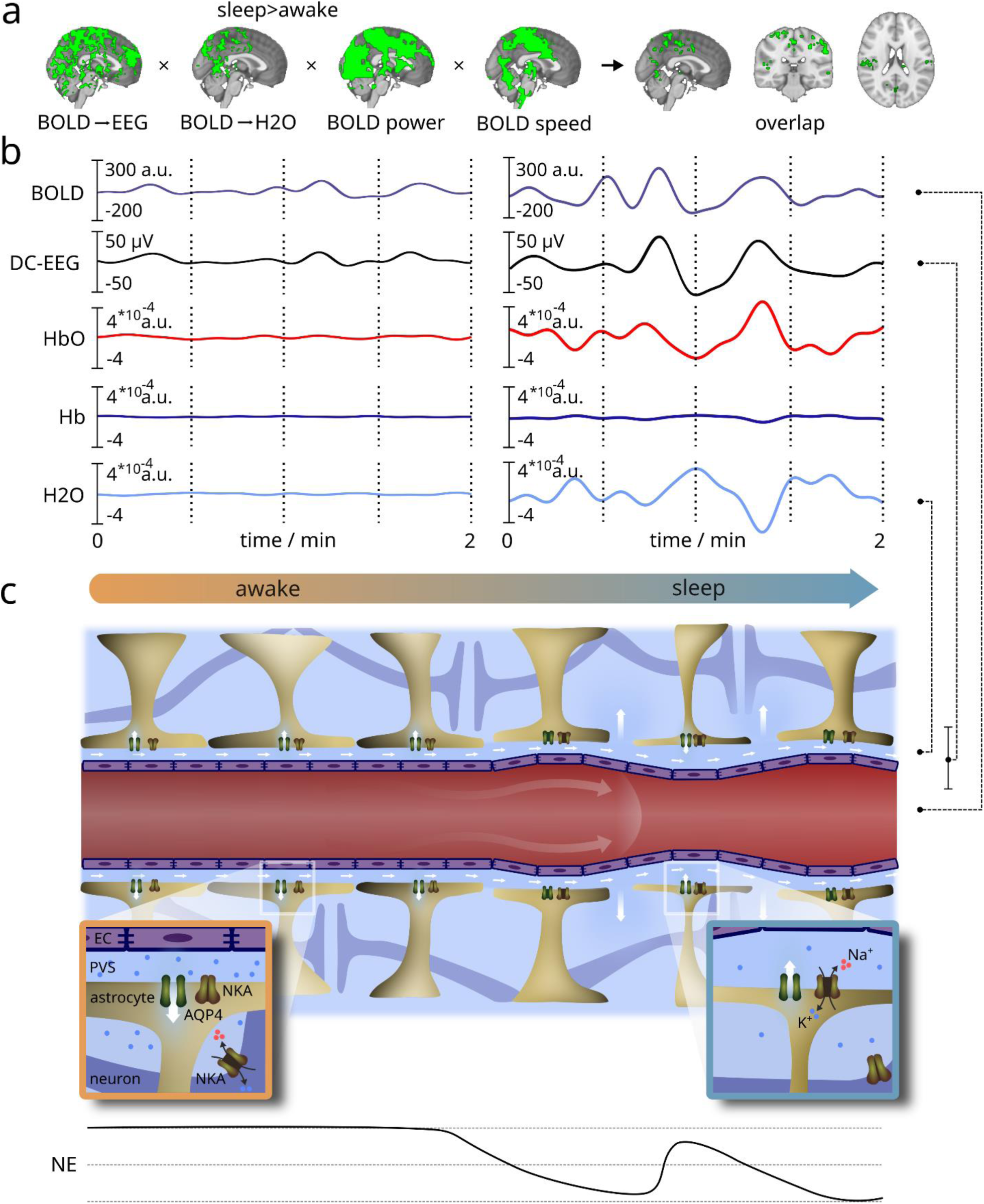
(a) Multimodal evidence of spatial overlap in primary sensory and motor areas, showing significant increases in sleep induced infra-slow (<0.1 Hz) pulsation power, vasomotor wave flow speeds^30^, and vasomotor-driven changes in both water dynamics and electrophysiological brain activity. (b) Concurrent infra-slow signals, capturing blood oxygen level-dependent (BOLD) signals, electroencephalography (DC-EEG), and functional near-infrared spectroscopy (fNIRS) during wakefulness and sleep. BOLD-calibrated, synchronous fNIRS signals indicate that the water (H₂O) signal is anti-correlated with arterial HbO₂ and BOLD signal^4,5,28^. Changes in venous Hb concentration are less pronounced, consistent with animal studies suggesting that BOLD signal generation is driven more by speed than by volume changes in the venous compartment^31^. (c) A theoretical illustration of regional electro-, hydro-, and hemodynamic interactions in wakefulness and NREM sleep. During NREM sleep, reduced orexin levels induce slow oscillations in norepinephrine (NE) levels^4,32^ and extracellular potassium [K+] concentration^33^. These oscillations drive vasomotor waves detected by both BOLD and oxygenated hemoglobin (HbO₂) signals^12,13,31^, which are accompanied by anti-correlated H₂O waves, as in recent animal findings^4,9^. Low NE levels reduce astrocytic volume by decreasing Na⁺/K⁺-ATPase (NAK) activity^2,4^. Pulsatile vasomotor waves cause oscillations in astrocytic endfeet and perivascular volumes^9^. These oscillations facilitate hydrodynamic interstitial/cerebrospinal fluid (I/CSF) exchange via enlarged inter-astrocytic gaps formed by astrocytic shrinkage. This mechanism provides in theory a generation mechanism for the slow but high-voltage infra-slow oscillations in EEG. Finally, the NE oscillations drive an important, increasing effect on neuronal NAK-channel activity that is opposed to the astrocytes effect closer to perivascular space. This opposing effect can explain the previously detected electro-osmotic potential difference increasing the I/CSF exchange over glia limitans explaining partially also the high voltage DC-EEG potential^25^.

## Discussion

In this study, we used a non-invasive multimodal neuroimaging setup to investigate the coupling of infra-slow (<0.1 Hz) brain oscillations of human brain including vasomotion (BOLDMREG), electrophysiological signals (DC-EEG), and CSF fluctuations (fNIRS) across the brain of healthy volunteers. During the awake state, low-power infra-slow electrophysiological potential and CSF oscillations both predicted vasomotor BOLD waves, consistent with previous findings on neurovascular coupling lag structures^31,34^. In contrast, during sleep, the amplitude dynamics increased along with a reversal in prediction patterns: the presumably NE-driven increases in vasomotor waves then began to coordinate electrophysiological activity and water dynamics, notably within parasagittal frontoparietal and temporal cortical areas. Notably, these regions also demonstrated increased speed and power of BOLD vasomotor waves during NREM sleep. In light of recent literature, our findings suggest that NE-induced vasomotor waves could propel CSF flow and further modulate electrophysiological oscillations of the cerebral cortex in human slow-wave sleep^1,4^.

### Increased pulsation powers are linked to sleep

We observed a significant increase in infra-slow BOLD power during sleep, consistent with previous findings^10–12,14,35^. Propagating infra-slow vasomotor waves in the BOLD signal^36–39^ reflect reciprocal oscillations in blood and CSF volume^4,5,28,36^. The sleep-regulating neuropeptide orexin, controls arousal levels by modulating LC activity ^40^, which is elevated in the awake state, resulting in increased cortical NE levels, thereby suppressing vasomotor waves. With type I narcolepsy - characterized by a specific depletion of orexin – the vasomotor waves are stronger in comparison to healthy controls^32^, supporting the role of orexin in regulating the NE-driven vasomotion.

Vasomotor BOLD waves have been related to autonomous nervous system activity in humans^41^ . Invasive animal experiments show that vasomotor waves are strongly linked to LC-generated slow fluctuations in extracellular NE levels^1,20^. Meanwhile, the T2* weighted fMRI signal of the brain cortex and blood vessels is anti-correlated with CSF pulsations in the ventricles^5,6,36^. Thus, NE oscillations cause anticorrelated blood and CSF oscillations; the descending phase of NE activity also correlates with the occurrence of sleep spindles, and enhanced glymphatic clearance of tracers from the brain^1,4,20^. Previous work has also shown that vasomotor waves all drive injected CSF tracers within arterial wall structures^42^.

Moreover, we detected increased power in infra-slow DC-EEG fluctuations during sleep, consistent with previous studies^16,17^. The mechanisms of the DC-EEG infra-slow oscillations have remained unknown. In sleep, the oscillating low levels of NE increase plasma membrane Na/K-ATPase activity in astrocytes located close to the BBB, while reducing its activity in neurons located further away^33,43^. These opposing responses of astrocytic and neuronal Na/K-ATPase activity could potentially generate an inter-cellular electrophysiological potential and consequently osmotic differences, which could drive local hydrodynamic flow due to osmoelectric effect^4,25,33^. During natural sleep, the thickness of astrocytic endfeet at the BBB, and perivascular space volume oscillates, which could together facilitate the hydrodynamic electrolyte oscillations over the BBB *glia limitans* in extending further into perivascular spaces via widened inter-astrocytic gaps^9,17,19^. These NE-driven oscillatory mechanisms could underlie the increased power of DC-EEG oscillations during reduced neuronal activity in sleep, as well as the increased synchrony between DC-EEG and neuronal activity during sleep^17–19^.

### Dynamics between electric, hydrodynamic and blood flow changes in awake human brain

Standard correlation metrics or lag analyses often fail to infer accurately the directed interactions, especially in the presence of non-linear interactions. Such approaches are susceptible to detecting spurious connections due to shared signal sources or artifacts^44,45^. To address this limitation, we employed phase transfer entropy (TE) - a non-linear extension of Granger causality - to accurately quantify putative causal interactions among physiological signals^29^.

In functional hyperemia, neural activation is followed by an increase in local blood flow, which is traditionally thought to enhance the supply of oxygen and glucose. However, recent findings suggest that synchronized neuronal activation also facilitates glymphatic solute transport and clearance^25,46–48^. In the activated areas, increased arterial pulsations typically occur 1.3 sec prior to the neuronally-coupled BOLD vasomotion^49^. As increased cardiovascular pulsations of local arteries may also increase glymphatic solute transport^50^, the vasodilatory pulsations could potentially serve as an additive mechanism to accommodate increased metabolic demands in addition to the osmoelectric neural activity^25,47,48^.

Consistent with this proposition, coupling patterns in awake brain revealed a sequential drive pattern in which electrophysiological activity predicted cortical water movement and CBF: DC-EEG → H2O → BLOOD, similarly as demonstrated in mice^4^. Water movement follows the neuronal [K+] release that is buffered by astrocytic of K^+^IR4/5 channels, with passive fluid flow further facilitated by neighboring astrocytic AQP4 water channels^51^. The local [K+] increases are also sensed by capillary endothelia K^+^IR4/5-channels, which may facilitate upstream vasomotor dilatations via inward hyperpolarizing waves mediated by gap junctions along blood vessels^52^. The occurrence of inward waves triggered by local [K+] increases offers a direct mechanism whereby NE effects on ion channels drive vascular vasomotor waves. Taken together, we find that, CSF volume changes and vasomotion both closely follow the electrophysiological activity in awake brain, likely playing a role in neuro-metabolic coupling to support functional increases in neuronal activity.

### Reversed brain fluid dynamics during sleep

The infra-slow vasomotor waves coordinated electrophysiological activity levels in DC-EEG and likewise cortical water volume fluctuations in the human cortex (see Fig. 3-4a). Large vasodilatory oscillations in intracranial blood volume, simultaneously captured here from BOLD signals and the arterial HbO2 signal in fNIRS (Fig 4b), were associated by opposing water concentration changes, consistent with the Monro-Kellie doctrine, namely that sum of volumes of brain, CSF, and intracerebral blood must be constant^4–6,9,28^.

We have demonstrated in an ultrasound-verified phantom experiment that optical flow analytics of MREG precisely measure water flow ^30^. Our previous human MREG results showed that the intracranial CSF propagation speed^30^ and power^12^ of vasomotor BOLD waves increase in sleep, particular in the primary sensory and motor regions, indicating an acceleration of brain water movement (see Fig. 4, supplementary Fig 1). Notably, neuronal slow delta activity also increases in same regions, which previous animal experimentation has linked to enhanced CSF solute transport ^12,53^. Furthermore, the infra-slow electrophysiological drive of brain water appears to increase in parietal areas during sleep (see Fig. 3 f,g), suggesting that brain water passively follows electrophysiological changes via osmoelectric effects, mediated through inter-astrocytic clefts and AQP4 channels^25^.

These findings, combined with the recent literature on increased hydrodynamic interstitial fluid flow via widened BBB *glia limitans* gaps, underscore the need for future studies to investigate these mechanisms at the cellular level, preferably using minimally invasive *in vivo* approaches that preserve the brain’s sensitive electro/hydrodynamics. Notably, the reversal of coupling patterns in human sleep observed in cortex did not occur in the deeper subcortical brain structures and white matter, where BOLD changes continued to follow DC-EEG and CSF volume fluctuations. This structural and functional-anatomic distinction aligns with previously observed differences in solute transport mechanisms between the cortex compared with deep brain structures, which showed slower tracer kinetics compared to the more efficient solute clearance in the cortical areas^54^.

## Materials and Methods

### Experimental setup

This study was approved by Regional Ethics Committee of the Northern Ostrobothnia Hospital District. We obtained written informed consent from all participants in accordance with the Declaration of Helsinki. The datasets utilized in this study were originally gathered as part of our prior investigation^12,13,17^ involving 24 healthy controls (13 females, 11 males). Participants underwent scanning twice across separate sessions: once during wakefulness and once during sleep. To increase sleep pressure, and thus enable faster onset of the sleep recordings, 13 subjects underwent one night of sleep deprivation, which was monitored with Oura rings (Oura Health Oy). Our study employed a multimodal neuroimaging setup^55^ to capture simultaneous hemodynamic, electrical, and physiological information. We used an MRI scanner to capture functional ultra-fast MREG and structural image series. Spontaneous brain activity was recorded with EEG, which was further used to derive sleep state-specific information. Sleep scoring was performed in 30-second epochs by experienced neurophysiologists according to AASM criteria (J.P., M.K.). Functional NIRS was used to measure changes in macroscopic water concentrations. All modalities were synchronized with the MRI scanner’s optical timing pulse. Calculations were performed using MATLAB (v.R2023b, MathWorks).

### MRI Acquisition and Preprocessing

Functional and structural images were acquired using a Siemens MAGNETOM Skyra 3T scanner equipped with a 32-channel head coil. For structural 3D MPRAGE scanning (TR=1900 ms, TE= 2.49 ms, TI=900 ms, FA=9°, FOV=240 mm), we used a slice thickness of 0.9 mm. Functional imaging was performed using an ultrafast MREG-sequence (TR=100 ms, TE=36 ms, FA=5°, FOV=192 mm), which employs k-space undersampling to reach a sampling frequency of 10 Hz and voxel size of 3 mm^56^. We set a crusher gradient to 0.1, which is optimized for detecting physiological signal sources, while mitigating slow drifts and stimulated echoes.

Reconstruction of MREG images involved the utilization of L2-Tikhonov regularization, wherein a lambda value of 0.1 was determined using the L-curve method^57^. Dynamic off-resonance correction in k-space was implemented to reduce B0-field artifacts and mitigate respiratory motion. Image preprocessing followed the standardized FSL (Functional Magnetic Resonance Imaging of the Brain’s software library) preprocessing pipeline^58^. A high-pass filter was used with a cut-off frequency of 0.008 Hz. Subsequently, we performed motion correction followed by brain extraction. To eliminate artifactual spikes, datasets were de-spiked^59^. The structural 3D MPRAGE images were used in the registration of functional datasets into the MNI152 standard space. Continuous two-minute segments of wakefulness and NREM-1/2 sleep were identified using sleep scores^13^, and filtered to the infra-slow frequency range (0.01-0.08 Hz).

### EEG Acquisition and Preprocessing

EEG recordings were acquired using a GES 400 (Magstim EGI) system, which included a direct current-coupled amplifier (Net Amps 400), and a high-density 256-electrode system (HydroCel Geodesic Sensor MR net) with the electrode ‘Cz’ serving as the reference channel. The recordings were conducted at a sampling rate of 1 kHz, except for three sleep and five awake subjects in whom the experimentalists had inadvertently selected 250 Hz. Prior to measurements, we performed a visual inspection of signal quality and electrode impedances.

The initial steps in data processing involved the removal of gradient and ballistocardiographic artifacts through template subtraction^60,61^, Brain Vision Analyzer v.2.1, Brain Products). Subsequently, bad channels were excluded based on the following criteria: standard deviation exceeding 2000 µV, average correlation with neighboring electrodes falling below 0.1, or electrode impedance surpassing 1 MΩ. Upon identifying bad channels, spherical interpolation was used to replace the removed channels^62^. EEG signals were decimated to 10 Hz to align with the sampling frequency of other modalities. A zero-phase bandpass filter (0.01-0.08 Hz) was employed using mirrored signals to eliminate filtering edge effects and phase distortions. Finally, measurements were segmented into sleep state-specific epochs.

### NIRS Acquisition and Preprocessing

Our functional NIRS device^28^ utilized a frequency-coding technique in which the emitted light was modulated at specific frequencies for each wavelength. High-power LEDs generated monochromatic light at wavelengths of 690, 830, and 980 nm, facilitating measurements of Hb, HbO, and H2O, respectively. Specifically, we selected the 980 nm wavelength for its high absorbance of water, ensuring high sensitivity in water dynamics measurement. At the receiver optode, the light was demodulated again with the corresponding frequencies. To maintain consistent optode positioning, we separated the source and receiver optodes on both sides of the EEG electrode ‘Fp1’ at a fixed distance of 3 cm, thereby ensuring a constant distance between measurements, while enabling near-infrared light to reach the underlying cerebral cortex^28^.

Initially, NIRS transmittance values were used to compute HbO, HbR, and water concentrations following the modified Beer-Lambert law^28,63^. NIRS concentration signals were decimated to 10 Hz to match the sampling rates of all three modalities. Subsequently, we used a zero-phase recursive bandpass filter (0.01-0.08 Hz) using the mirroring method to isolate the infra-slow frequencies. Finally, we extracted waking and sleep state-specific epochs over the same time points as those for EEG and MREG.

### Estimation of spectral properties

We employed complex Morlet wavelets in wavelet convolution to conduct time-frequency spectral analysis (see Fig. 2a,b). These wavelets consist of sine waves at various frequencies tapered by a Gaussian window, thereby providing temporal specificity. Convolving time series with each frequency’s wavelet transforms the data into frequency-specific power by computing the squared magnitude of the convolution results. Unlike rectangular windowing, the Gaussian window reduces ripple effects typically associated with sharp edges in the kernel signal. We chose wavelet convolution due to its ability to focus on time-domain changes, its computational efficiency, and its adherence to the assumption of stationarity, contrary to conventional spectral methods^64^. We applied a logarithmic frequency range extending from 0.01 Hz to 5 Hz with 50 steps, having selected the 5 Hz upper limit as it represents the Nyquist frequency of the MREG recordings. To mitigate edge effects that could potentially contaminate the time series, particularly in low-frequency filtering, we implemented the mirroring technique. The number of wavelet cycles (*N*=8) was held constant, regulating the trade-off between frequency and temporal precision. Given that we had nearly 70,000 spatially-correlating signals in each MREG dataset, we employed a simple random sampling technique without replacement to reduce computational cost. Here, we used 5% (3,405 voxels) of all voxels, ensuring that each brain voxel had an equal probability of being chosen. Finally, we calculated power in the frequency range of 0.01-0.08 Hz using integral approximation and rectangular windowing.

### Inferring directed coupling patterns

Transfer entropy (TE) is a dynamic and directed measure of predictive information^29,65^. Essentially, TE represents the average information from a source signal that aids in predicting the next value of the target concerning its past. Conditioning in TE can have two opposing effects, either reducing TE by eliminating redundant information between the source and target past, or increasing TE by including synergistic information^44,66,67^. We calculated TE in the phase domain, thereby mitigating effects of linear mixing and noise^29^. To utilize the phase-based TE, we took the infra-slow time series and applied a Hilbert transform to compute the analytical signals *z*_*ISF*_(*n*). From these analytical signals, we extracted the instantaneous phases as *θ*_ISF_ = *arg*(*z*_ISF_(*n*)). In calculating TE (see Fig. 3), we chose a discrete estimator to represent the state space, being the simplest form of estimators. If the source-target delay is set to maximize information transfer, it aligns with the actual causal delay under simple conditions. However, predictive information can be accurately estimated over a wide range of lags^29^. Here, we assumed a constant delay of one cycle, in consideration of the computational cost of identifying the maximal information transfer. We calculated pairwise TE for each MREG voxel, including all EEG electrodes and water concentration signals. To infer the net direction of the information transfer, we took the difference as Δ*TE*(*x*, *y*) = *TE*(*x*, *y*) − *TE*(*y*, *x*).

### Vasomotor wave propagation speed

To investigate vasomotor wave propagation within the human brain, we applied optical flow analysis to the MREG data, focusing specifically on infra-slow oscillatory components associated with vasomotor activity. Optical flow analysis has been previously used for tracking motion within imaging data for different physiological pulsations including cardiac^68^ and respiratory pulsations^69^. By leveraging this method, we aimed to capture the spatiotemporal dynamics of vasomotor waves^30^. We quantified the velocity and spatial extent of vasomotor wave propagation, thereby characterizing how these waves propagate across cortical and subcortical regions.

Pulse-triggering was guided by identifying regions demonstrating the highest correlations with vasomotor activity, following established methodologies. Specifically, the posterior cingulate cortex emerged as the region most strongly associated with vasomotor oscillations, leading to its selection as the primary reference region for triggering vasomotor wave analysis. This approach provided a precise framework for delineating the onset and propagation of vasomotor oscillations within the brain’s functional architecture.

### Statistical analysis

We had not made *a priori* statistical power calculation to predetermine the sample size for this study, but used the material at hand. The alpha level was set to 0.05 for all statistical tests. We investigated whether infra-slow power differed between epochs of wakefulness and the combined sleep states (NREM-1/2). A Wilcoxon rank-sum test (two-tailed) was employed, with the null hypothesis being that awake and sleep epochs had equal infra-slow power (see Fig. 2b). The false discovery rate was controlled using FDR correction^70^ to account for multiple comparisons. To compare voxelwise group differences in TE, we conducted a randomization test with 5,000 permutations, shuffling the arousal state labels for awake; NREM-1, awake; NREM-2, and NREM-1; NREM-2 (see Fig. 3c, e). Here we used the threshold-free cluster enhancement method to address multiple comparisons. Additionally, to compare whole-brain average TE, we used a Wilcoxon rank-sum test (two-tailed), with the null hypothesis being that arousal states had equal median information transfer (see Fig. 3c, e, g). For comparison of BOLD velocity and speed maps between awake and sleep states we used randomization test along with threshold-free cluster enhancement^30^.

## Acknowledgments

This study was funded by Instrumentarium Science Foundation (T.V), Jane & Aatos Erkko Foundation 1,210043 (V.Ki.), Academy of Finland TERVA grants 1-2 314497, 335720 (V.Ki), VTR grants from Oulu University Hospital (V.Ki, V.Ko), The EU Joint Programme – Neurodegenerative Disease Research 2022-120 (V.Ki), Academy of Finland Grant 275342, 338599 (V.Ki), The Finnish Medical Foundation (V.Ki), Finnish Brain Foundation (V.Ki), Uniogs/MRC Oulu DP-grant (H.H), Emil Aaltosen Säätiö (H.H), Pohjois-Suomen Terveydenhuollon tukisäätiö (H.H, V.Ko). We thank CSC – IT Center for Science Ltd., Finland for providing high-performance computational resources used in this study. We further thank Hannu Kinnunen and Oura Health for their collaboration. We thank Prof. Paul Cumming of Bern University Hospital for comments on the manuscript.

## References

1. Kjaerby, C. et al. Memory-enhancing properties of sleep depend on the oscillatory amplitude of norepinephrine. Nat Neurosci 25, 1059–1070 (2022).

2. Xie, L. et al. Sleep drives metabolite clearance from the adult brain. Science 342, 373–377 (2013).

3. Osorio-Forero, A., Cherrad, N., Banterle, L., Fernandez, L. M. J. & Lüthi, A. When the Locus Coeruleus Speaks Up in Sleep: Recent Insights, Emerging Perspectives. Int J Mol Sci 23, 5028 (2022).

4. Hauglund, N. L. et al. Norepinephrine-mediated slow vasomotion drives glymphatic clearance during sleep. Cell (2025) doi:10.1016/j.cell.2024.11.027.

5. Borchardt, V. et al. Inverse correlation of fluctuations of cerebral blood and water concentrations in humans. The European Physical Journal Plus 136, 497 (2021).

6. Fultz, N. E. et al. Coupled electrophysiological, hemodynamic, and cerebrospinal fluid oscillations in human sleep. Science (1979) 366, 628–631 (2019).

7. Bork, P. A. R. et al. Astrocyte endfeet may theoretically act as valves to convert pressure oscillations to glymphatic flow. J R Soc Interface 20, (2023).

8. Gan, Y. et al. Perivascular pumping of cerebrospinal fluid in the brain with a valve mechanism. J R Soc Interface 20, (2023).

9. Bojarskaite, L. et al. Sleep cycle-dependent vascular dynamics in male mice and the predicted effects on perivascular cerebrospinal fluid flow and solute transport. Nat Commun 14, 953 (2023).

10. Chang, C. et al. Tracking brain arousal fluctuations with fMRI. Proceedings of the National Academy of Sciences 113, 4518–4523 (2016).

11. Fukunaga, M. et al. Large-amplitude, spatially correlated fluctuations in BOLD fMRI signals during extended rest and early sleep stages. Magn Reson Imaging 24, 979–992 (2006).

12. Helakari, H. et al. Human NREM Sleep Promotes Brain-Wide Vasomotor and Respiratory Pulsations. The Journal of Neuroscience 42, 2503–2515 (2022).

13. Helakari, H. et al. Effect of sleep deprivation and NREM sleep stage on physiological brain pulsations. Front Neurosci 17, (2023).

14. Horovitz, S. G. et al. Low frequency BOLD fluctuations during resting wakefulness and light sleep: A simultaneous EEG-fMRI study. Hum Brain Mapp 29, 671–682 (2008).

15. Liu, Z., Fukunaga, M., de Zwart, J. A. & Duyn, J. H. Large-scale spontaneous fluctuations and correlations in brain electrical activity observed with magnetoencephalography. Neuroimage 51, 102–111 (2010).

16. Marshall, L., Molle, M., Michaelsen, S., Fehm, H. L. & Born, J. Slow potential shifts at sleep--wake transitions and shifts between NREM and REM sleep. Sleep 19, 145–151 (1996).

17. Väyrynen, T. et al. Infra-slow fluctuations in cortical potentials and respiration drive fast cortical EEG rhythms in sleeping and waking states. Clinical Neurophysiology 156, 207–219 (2023).

18. Monto, S., Palva, S., Voipio, J. & Palva, J. M. Very Slow EEG Fluctuations Predict the Dynamics of Stimulus Detection and Oscillation Amplitudes in Humans. The Journal of Neuroscience 28, 8268–8272 (2008).

19. Vanhatalo, S. et al. Infraslow oscillations modulate excitability and interictal epileptic activity in the human cortex during sleep. Proc Natl Acad Sci U S A 101, 5053–5057 (2004).

20. Osorio-Forero, A. et al. Noradrenergic circuit control of non-REM sleep substates. Current Biology 31, 5009–5023.e7 (2021).

21. Antony, J. W. et al. Sleep Spindle Refractoriness Segregates Periods of Memory Reactivation. Current Biology 28, 1736–1743.e4 (2018).

22. Kiviniemi, V. et al. Real-time monitoring of human blood-brain barrier disruption. PLoS One 12, e0174072 (2017).

23. Voipio, J., Tallgren, P., Heinonen, E., Vanhatalo, S. & Kaila, K. Millivolt-Scale DC Shifts in the Human Scalp EEG: Evidence for a Nonneuronal Generator. J Neurophysiol 89, 2208–2214 (2003).

24. Besson, J. M. et al. Correlations of brain d-c shifts with changes in cerebral blood flow. Am J Physiol 218, 284–291 (1970).

25. Jiang-Xie, L.-F. et al. Neuronal dynamics direct cerebrospinal fluid perfusion and brain clearance. Nature 627, 157–164 (2024).

26. Chong, P. L. H., Garic, D., Shen, M. D., Lundgaard, I. & Schwichtenberg, A. J. Sleep, cerebrospinal fluid, and the glymphatic system: A systematic review. Sleep Med Rev 61, 101572 (2022).

27. Huotari, N. et al. Sampling Rate Effects on Resting State fMRI Metrics. Front Neurosci 13, (2019).

28. Myllylä, T. et al. Assessment of the dynamics of human glymphatic system by near-infrared spectroscopy. J Biophotonics 11, (2018).

29. Lobier, M., Siebenhühner, F., Palva, S. & Palva, J. M. Phase transfer entropy: A novel phase-based measure for directed connectivity in networks coupled by oscillatory interactions. Neuroimage 85, 853–872 (2014).

30. Elabasy, A. et al. Sleep increases propagation speed of physiological brain pulsations. bioRxiv 2025.01.09.632145 (2025) doi:10.1101/2025.01.09.632145.

31. Hillman, E. M. C. Coupling Mechanism and Significance of the BOLD Signal: A Status Report. Annu Rev Neurosci 37, 161–181 (2014).

32. Järvelä, M. et al. Increased very low frequency pulsations and decreased cardiorespiratory pulsations suggest altered brain clearance in narcolepsy. Communications Medicine 2, 122 (2022).

33. Li, B. et al. Anti-seizure effects of norepinephrine-induced free fatty acid release. Cell Metab 37, 223–238.e5 (2025).

34. Logothetis, N. K. et al. The effects of electrical microstimulation on cortical signal propagation. Nat Neurosci 13, 1283–1291 (2010).

35. Liu, X. et al. Subcortical evidence for a contribution of arousal to fMRI studies of brain activity. Nat Commun 9, 395 (2018).

36. Kiviniemi, V. et al. Ultra-fast magnetic resonance encephalography of physiological brain activity – Glymphatic pulsation mechanisms? Journal of Cerebral Blood Flow & Metabolism 36, 1033–1045 (2016).

37. Rayshubskiy, A. et al. Direct, intraoperative observation of ∼ 0.1 Hz hemodynamic oscillations in awake human cortex: Implications for fMRI. Neuroimage 87, 323–331 (2014).

38. Wang, H. H. et al. Physiological noise in MR images: An indicator of the tissue response to ischemia? Journal of Magnetic Resonance Imaging 27, 866–871 (2008).

39. Bolt, T. et al. A parsimonious description of global functional brain organization in three spatiotemporal patterns. Nat Neurosci 25, 1093–1103 (2022).

40. Horvath, T. L. et al. Hypocretin (orexin) activation and synaptic innervation of the locus coeruleus noradrenergic system. J Comp Neurol 415, 145–59 (1999).

41. Picchioni, D. et al. Autonomic arousals contribute to brain fluid pulsations during sleep. Neuroimage 249, 118888 (2022).

42. van Veluw, S. J. et al. Vasomotion as a Driving Force for Paravascular Clearance in the Awake Mouse Brain. Neuron 105, 549–561.e5 (2020).

43. Wotton, C. A., Cross, C. D. & Bekar, L. K. Serotonin, norepinephrine, and acetylcholine differentially affect astrocytic potassium clearance to modulate somatosensory signaling in male mice. J Neurosci Res 98, 964–977 (2020).

44. Bossomaier, T., Barnett, L., Harré, M. & Lizier, J. T. An Introduction to Transfer Entropy: Information Flow in Complex Systems. (Springer International Publishing, 2016). 10.1007/978-3-319-43222-9.

45. Murari, A. et al. On the Use of Transfer Entropy to Investigate the Time Horizon of Causal Influences between Signals. Entropy (Basel) 20, (2018).

46. Holstein-Rønsbo, S. et al. Glymphatic influx and clearance are accelerated by neurovascular coupling. Nat Neurosci 26, 1042–1053 (2023).

47. Murdock, M. H. et al. Multisensory gamma stimulation promotes glymphatic clearance of amyloid. Nature 627, 149–156 (2024).

48. Kim, E., Van Reet, J. & Yoo, S.-S. Cerebrospinal fluid solute transport associated with sensorimotor brain activity in rodents. Sci Rep 13, 17002 (2023).

49. Huotari, N. et al. Cardiovascular Pulsatility Increases in Visual Cortex Before Blood Oxygen Level Dependent Response During Stimulus. Front Neurosci 16, (2022).

50. Mestre, H. et al. Flow of cerebrospinal fluid is driven by arterial pulsations and is reduced in hypertension. Nat Commun 9, 4878 (2018).

51. Nagelhus, E. A., Mathiisen, T. M. & Ottersen, O. P. Aquaporin-4 in the central nervous system: Cellular and subcellular distribution and coexpression with KIR4.1. Neuroscience 129, 905–913 (2004).

52. Longden, T. A. et al. Capillary K+-sensing initiates retrograde hyperpolarization to increase local cerebral blood flow. Nat Neurosci 20, 717–726 (2017).

53. Ju, Y.-E. S. et al. Slow wave sleep disruption increases cerebrospinal fluid amyloid-β levels. Brain 140, 2104–2111 (2017).

54. Ringstad, G. & Eide, P. K. Cerebrospinal fluid tracer efflux to parasagittal dura in humans. Nat Commun 11, 354 (2020).

55. Korhonen, V. et al. Synchronous Multiscale Neuroimaging Environment for Critically Sampled Physiological Analysis of Brain Function: Hepta-Scan Concept. Brain Connect 4, 677–689 (2014).

56. Assländer, J. et al. Single shot whole brain imaging using spherical stack of spirals trajectories. Neuroimage 73, 59–70 (2013).

57. Hugger, T. et al. Fast Undersampled Functional Magnetic Resonance Imaging Using Nonlinear Regularized Parallel Image Reconstruction. PLoS One 6, e28822 (2011).

58. Jenkinson, M., Beckmann, C. F., Behrens, T. E. J., Woolrich, M. W. & Smith, S. M. FSL. Neuroimage 62, 782–790 (2012).

59. Cox, R. W. AFNI: Software for Analysis and Visualization of Functional Magnetic Resonance Neuroimages. Computers and Biomedical Research 29, 162–173 (1996).

60. Allen, P. J., Josephs, O. & Turner, R. A method for removing imaging artifact from continuous EEG recorded during functional MRI. Neuroimage 12, 230–239 (2000).

61. Allen, P. J., Polizzi, G., Krakow, K., Fish, D. R. & Lemieux, L. Identification of EEG events in the MR scanner: the problem of pulse artifact and a method for its subtraction. Neuroimage 8, 229–239 (1998).

62. Oostenveld, R., Fries, P., Maris, E. & Schoffelen, J.-M. FieldTrip: Open Source Software for Advanced Analysis of MEG, EEG, and Invasive Electrophysiological Data. Comput Intell Neurosci 2011, 1–9 (2011).

63. Cope, M. & Delpy, D. T. System for long-term measurement of cerebral blood and tissue oxygenation on newborn infants by near infra-red transillumination. Med Biol Eng Comput 26, 289–294 (1988).

64. Cohen, M. X. Analyzing Neural Time Series Data. (The MIT Press, 2014). doi:10.7551/mitpress/9609.001.0001.

65. Schreiber, T. Measuring Information Transfer. Phys Rev Lett 85, 461–464 (2000).

66. Lizier, J. T. & Prokopenko, M. Differentiating information transfer and causal effect. Eur Phys J B 73, 605–615 (2010).

67. Wibral, M. et al. Measuring Information-Transfer Delays. PLoS One 8, e55809 (2013).

68. Rajna, Z. et al. Cardiovascular brain impulses in Alzheimer’s disease. Brain 144, 2214–2226 (2021).

69. Elabasy, A. et al. Respiratory brain impulse propagation in focal epilepsy. Sci Rep 13, 5222 (2023).

70. Benjamini, Y. & Hochberg, Y. Controlling the False Discovery Rate: A Practical and Powerful Approach to Multiple Testing. Journal of the Royal Statistical Society. Series B (Methodological) 57, 289–300 (1995).

