## Supplementary Figure 1 for "Vasomotor Pulsation Becomes a Driver of Cerebrospinal Fluid and Electrophysiological Dynamics in Sleeping Human Brain"

### Supplementary information

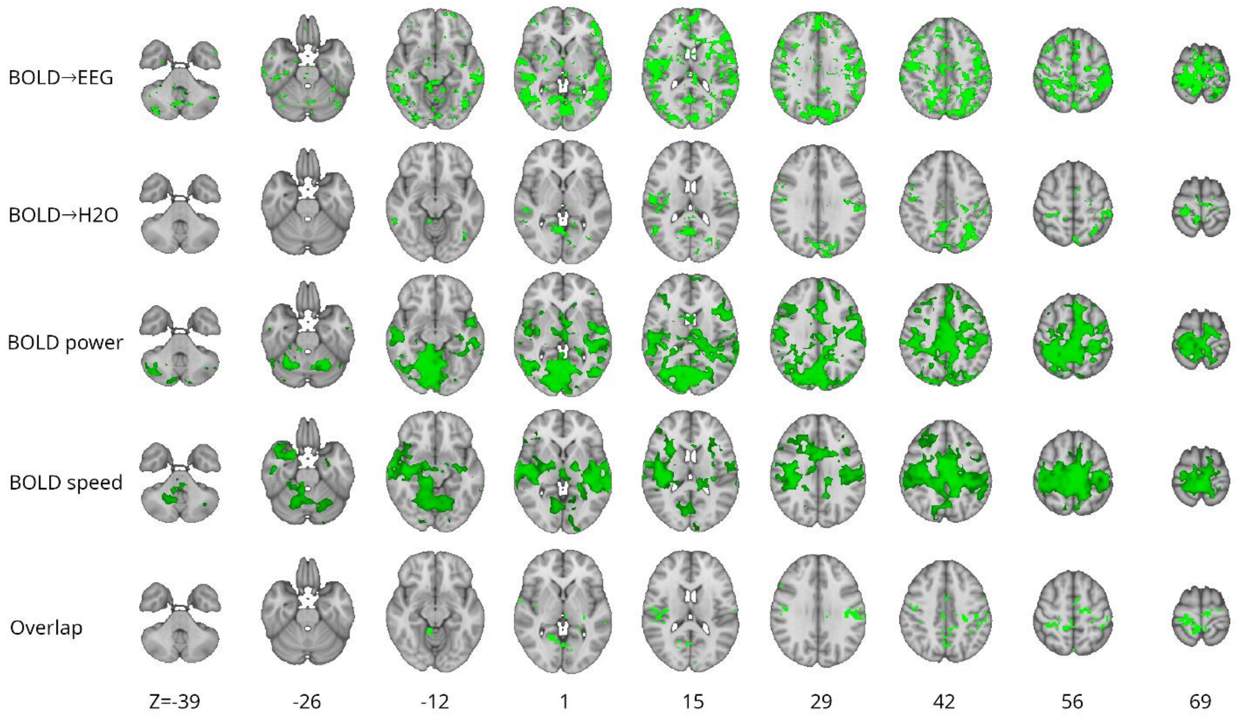


**Supplementary figure 1**. Statistically significant (sleep>awake, $p<0.05$) brain regions for sleep related BOLD signal prediction of infra-slow EEG and $H_2O$ oscillations, BOLD power and BOLD vasomotor wave propagation speed with both NREM1 & NREM2 combined in each analysis. The bottom row shows the overlap between these metrics. The coordinate system is represented in MNI.
